# First hybrid complete genome of *Aeromonas veronii* reveals chromosome-mediated novel structural variant *mcr*-*3.19* from human clinical specimen

**DOI:** 10.1101/576421

**Authors:** Naveen Kumar Devanga Ragupathi, Dhiviya Prabaa Muthuirulandi Sethuvel, Shalini Anandan, Divya Murugan, Kalaiarasi Asokan, Ramya Gajaraj Neethi Mohan, Karthick Vasudevan, Thirumal Kumar D, George Priya Doss C, Balaji Veeraraghavan

**Affiliations:** Department of Clinical Microbiology, Christian Medical College, Vellore – 632004, India; School of Bio Sciences and Technology, Vellore Institute of Technology, Vellore – 632014

**Author notes:** Corresponding author Dr. Balaji Veeraraghavan, Professor, Department of Clinical Microbiology Christian Medical College, Vellore – 632004. These authors have contributed equally to this work (Naveen Kumar Devanga Ragupathi, Dhiviya Prabaa Muthuirulandi Sethuvel).

**Keywords:** *mcr-3*, colistin, *Aeromonas*, ISAs*18*, *bla*OXA-12, *bla*CEPH-A3

## Abstract

**Background:** Recent findings substantiate the origin of plasmid-mediated colistin resistance gene *mcr-3* from Aeromonads. The present study aimed to screen the plasmid-mediated colistin resistance among 30 clinical multidrug resistant (MDR) *Aeromonas spp*.

**Results:** The presence of *mcr-1, mcr-2, mcr-3, and mcr-4* were screened by PCR, which revealed *mcr-3* in a colistin susceptible isolate (FC951). All other isolates were negative for *mcr* genes. Sequencing of FC951 revealed that *mcr-3* (*mcr-3.19*) identified was different from previously reported variants and had 95.62 and 95.28% nucleotide similarity with *mcr-3.3* and *mcr-3.10* gene. A hybrid assembly using IonTorrent and MinION reads revealed structural genetic information of *mcr-3.19* with an insertion of ISAs*18* within the gene. Due to this, *mcr-3.19* was non-expressive which makes FC951 susceptible to colistin. Further, *in silico* sequence and protein structural analysis confirmed the new variant. To the best of our knowledge, this is the first report on novel *mcr-3* variant (*mcr-3.19*).

**Conclusions:** The significant role of *mcr*-like genes in different *Aeromonas* species remains unknown and needs additional investigation to understand the insights on colistin resistance mechanism.

## Background

*Aeromonas spp* are ubiquitous and were known to cause gastroenteritis, wound infections, and septicemia, commonly known as “Jack of all trades”. Aeromonads had been universally resistant to the penicillin group of antibiotics (penicillin, ampicillin, carbenicillin, and ticarcillin) and are generally susceptible to tetracyclines and quinolones [1]. Recently, increasing resistance to third-generation cephalosporins and carbapenems are reported [2, 3].

The recent finding of a plasmid-mediated colistin resistance genes (*mcr*) has attracted global attention. A study reported that the *mcr-3* gene identified in *E. coli* was found similar to the gene present in *Aeromonas* species and suggested that it might have originated from Aeromonads [4]. It should be noted that most of the *Aeromonas* species are susceptible to colistin, whereas *A. jandaei* and *A. hydrophila* were reported as intrinsically resistant to polymyxins [4]. However, the significant role of *mcr*-like genes in different *Aeromonas* species is not clearly known and needs further investigation.

This study examined the presence of plasmid-mediated colistin resistance among *Aeromonas spp* by PCR and structural analysis using next-generation sequencing. The obtained nucleotide sequences from experimental methods were translated into protein sequence, and the 3D structure was modeled using *in silico* approaches to understand the structural changes in the different variants of *mcr* genes.

## Results

### Antimicrobial susceptibility

The resistance profile for the studied *Aeromonas* isolates were given in Table 1. Colistin MIC was determined for all the isolates which showed 30% of the isolates were found resistant to colistin. The MIC was identified to be 0.5 µg/ml (susceptible) for the isolate positive for *mcr-3* gene.

**Table 1:** Antimicrobial susceptibility of the selected *Aeromonas spp*

| S. no. | Sample ID | Age/sex | Organism | Resistance Pattern (Disc Diffusion) | Colistin MIC (µg/ml) |
| --- | --- | --- | --- | --- | --- |
| 1 | FC3340 | 29M | *Aeromonas* spp | AMP-SXT-TAX | 4R |
| 2 | FC193 | 0M | *Aeromonas dhakensis* | AMP-IMI-MEM | ≥64R |
| 3 | FC199 | 76M | *Aeromonas hydrophila* | AMP | 1S |
| 4 | FC284 | 75M | *Aeromonas* spp | AMP-IMI-MEM | 16R |
| 5 | FC728 | 71M | *Aeromonas caviae* | AMP-TET-TAX-IMI-MEM-CIP | 0.5S |
| 6 | FC729 | 66M | *Aeromonas caviae* | AMP-TET-SXT-TAX-IMI-MEM-CIP | 0.5S |
| 7 | FC715 | 34F | *Aeromonas caviae* | AMP | >32R |
| 8 | FC850 | 27M | *Aeromonas veronii* | AMP-TET-SXT-IMI-MEM-CIP | 1S |
| 9 | *FC951 | 55M | *Aeromonas veronii* | AMP-TET-IMI-MEM | 0.5S |
| 10 | FC1239 | 21F | *Aeromonas dhakensis* | AMP-TAX-IMI-MEM | ≥64R |
| 11 | FC1520 | 45F | *Aeromonas hydrophila* | AMP | >32R |
| 12 | FC1169 | 0M | *Aeromonas hydrophila* | AMP | 2S |
| 13 | FC599 | 57F | *Aeromonas hydrophila* | AMP-SXT | 2S |
| 14 | FC814 | 55F | *Aeromonas hydrophila* | AMP-IMI | >32R |
| 15 | FC2457 | 0M | *Aeromonas hydrophila* | AMP-TET-SXT | 2S |
| 16 | FC538 | 58M | *Aeromonas hydrophila* | AMP-IMP | 2S |
| 17 | FC771 | 1M | *Aeromonas caviae* | AMP-TAX-CIP | 4R |
| 18 | FC578 | 26F | *Aeromonas caviae* | AMP | 1S |
| 19 | FC377 | 22F | *Aeromonas hydrophila* | AMP-SXT-IMI-MEM | 1S |
| 20 | FC1245 | 42F | *Aeromonas hydrophila* | AMP-IMI | 2S |
| 21 | FC788 | 0M | *Aeromonas hydrophila* | AMP-TAX-IMI | 8R |
| 22 | FC157 | 58M | *Aeromonas veronii* | AMP | 0.5S |
| 23 | FC390 | 9F | *Aeromonas caviae* | AMP | 1S |
| 24 | FC1411 | 5M | *Aeromonas hydrophila* | AMP-IMI | 2S |
| 25 | FC523 | 1M | *Aeromonas hydrophila* | AMP-TET | 1S |
| 26 | FC1999 | 54F | *Aeromonas hydrophila* | AMP-IMI-MEM | 0.5S |
| 27 | FC2051 | 28M | *Aeromonas caviae* | AMP | 1S |
| 28 | FC2019 | 60M | *Aeromonas caviae* | AMP | 1S |
| 29 | FC906 | 55M | *Aeromonas hydrophila* | AMP-TET | 0.5S |
| 30 | FC1435 | 73F | *Aeromonas hydrophila* | AMP | >32R |

\* Isolate sequenced, R – resistant, S – susceptible, AMP – ampicillin, TET – tetracycline, SXT – trimethoprim/sulfamethoxazole, CIP – ciprofloxacin, TAX – cefotaxime, IMI – imipenem, MEM - meropenem

### Screening of *mcr* genes

Of the 30 isolates screened for *mcr-1, mcr-2, mcr-3* and *mcr-4* genes, only one isolate (FC951) was positive for *mcr-3* gene.

### Next-generation sequencing

*A. veronii* (FC951) positive for *mcr-3* gene by PCR was sequenced using IonTorrent PGM. Analysis of *mcr-3* gene revealed only 95.6% identity against the reference sequences in the database (henceforth termed as *mcr-3.19*). Also, ISFinder revealed ISAs*18* belonging to IS*4* family next to *eptA* aka *mcr-3*.

However, the *mcr-3.19* gene was split in to two contigs (IonTorrent). Generally, the IonTorrent assembly is highly accurate, but the assembly had too many fragments. The long read sequencing of FC951 using MinION resulted a complete genome (a chromosome and a plasmid), yet with errors. Hybrid assembly using IonTorrent and MinION reads in Unicycler resulted a complete chromosomal contig for FC951 with increased accuracy and low errors. Interestingly, the analysis of complete genome revealed *mcr-3.19* gene integrated in the chromosome along with the insertion of ISAs*18* within *mcr-3.19*. The structure of genetic environment of *mcr-3.19* was given in Figure 1A.

**Figure 1:**
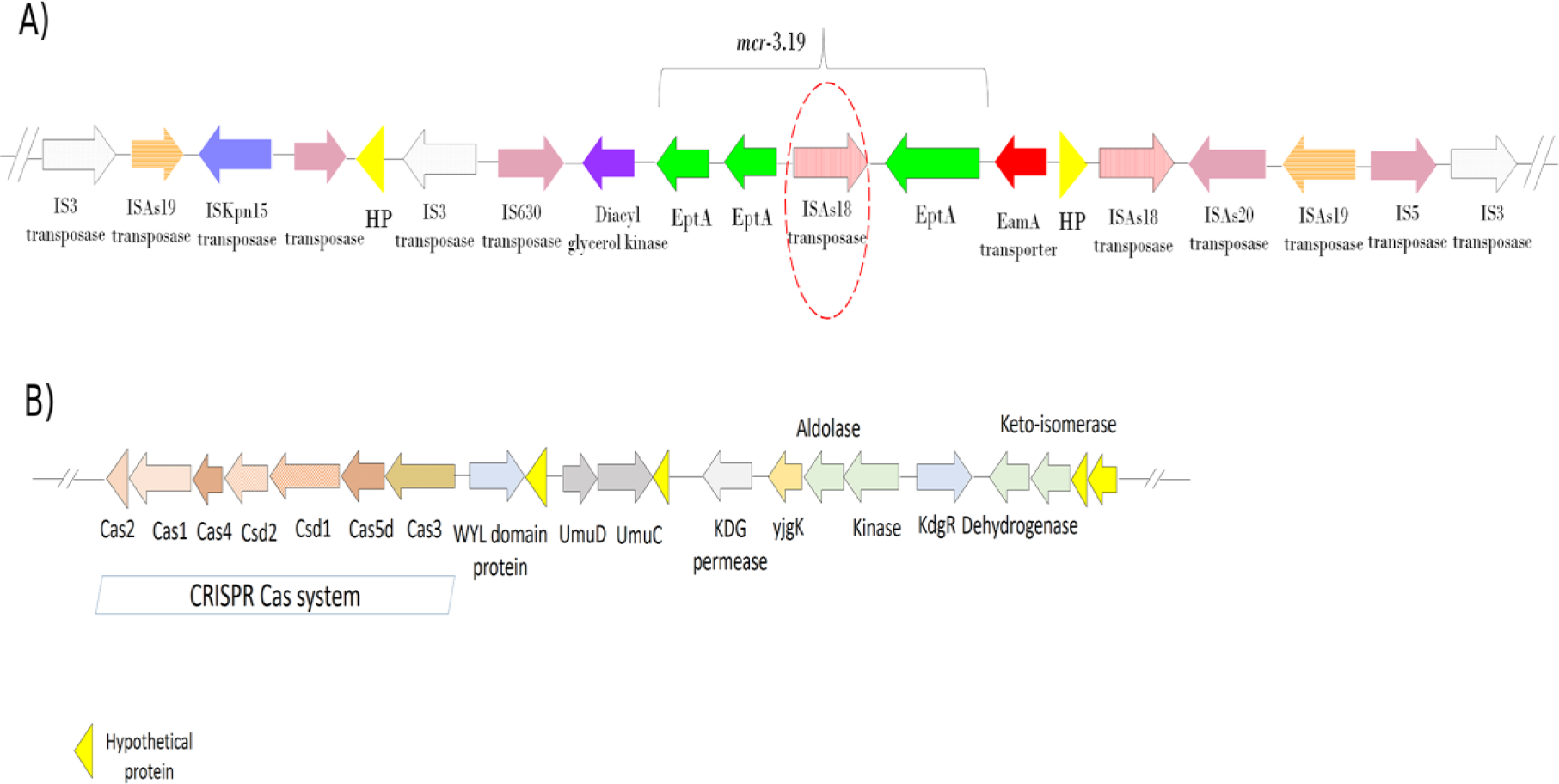
**A)** Genetic environment of *mcr-3.19* gene with an insertion of ISAs*18* transposase (1141 bp) leading to disruption of *mcr-3.19* functioning, **B)** CRISPR Cas system identified in FC951 and arrangements of Cas genes.

The Quast analysis depicted that N50 and N75 value of hybrid assembly to be 4660178, which is approximately 96% of the total assembly length. In addition, ANI is calculated using different assembly methods. It is evident that the closeness between IonTorrent and hybrid assembly is about 99.92%, which represent the high accuracy of hybrid assembly (Figure 2). The hybrid assembly generated 4.66-Mbp chromosome length single contig. In contrast, the IonTorrent-only assembly produced an assembly with more than 300 contigs and only 139 contigs >= 1000 bp. It was clear that hybrid assembly have its own advantage with improved accuracy and reduced error rate with genome completeness than MinION only or IonTorrent only assemblies.

**Figure 2:**
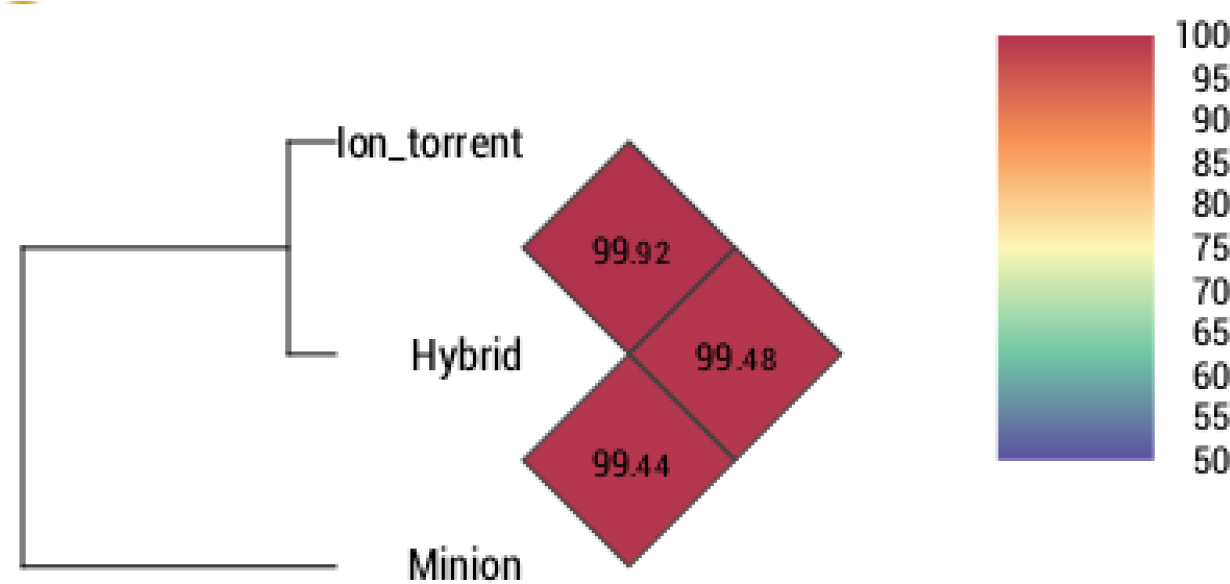
Representation of average nucleotide identities (ANI) using IonTorrent, MinION and hybrid assemblies.

Moreover, annotation of the extra-chromosomal sequences from MinION could not be designated as a complete plasmid. Though it showed 21% similarity with a previously reported *Xanthomonas citri* plasmid (CP020883.1).

Further analysis of resistance genes using ResFinder revealed the presence of *bla*OXA12 and *bla*CEPH-A3 genes. A CRISPR Cas system was identified in FC951 and arrangements of genes were as given in Figure 1B. Notably, sequence type of FC951 was identified to be novel, ST-515.

This complete genome project has been deposited at GenBank under the accession number CP032839 (Plasmid accession number CP032840).

### *In silico* sequence analysis

The *mcr-3* nucleotide and protein sequence identified in this study was compared with the previously reported variants *mcr-3.1* – *mcr-3.10* using BLAST search (KY924928.1, NMWW01000143.1, MF495680, NQCO01000074.1, MF489760, MF598076.1, MF598077.1, MF598078.1, MF598080.1 and MG214531) and identified to be a novel variant (*mcr-3.19*). The nucleotide percentage identity matrix for the *mcr* gene variants was as given in Table 2. From the analysis, *mcr-3.3* and *mcr-3.10* were found to be highly identical with 95.62 and 95.28 percentage respectively. The highest conserved amino acid region among the three variants (*mcr-3.3*, *mcr-3.10*, and *mcr-3.19*) were identified using Clustal Omega. The region of amino acids from LEU-359 to ILE-427 was found to be the largest conserved sequence among them.

**Table 2:**
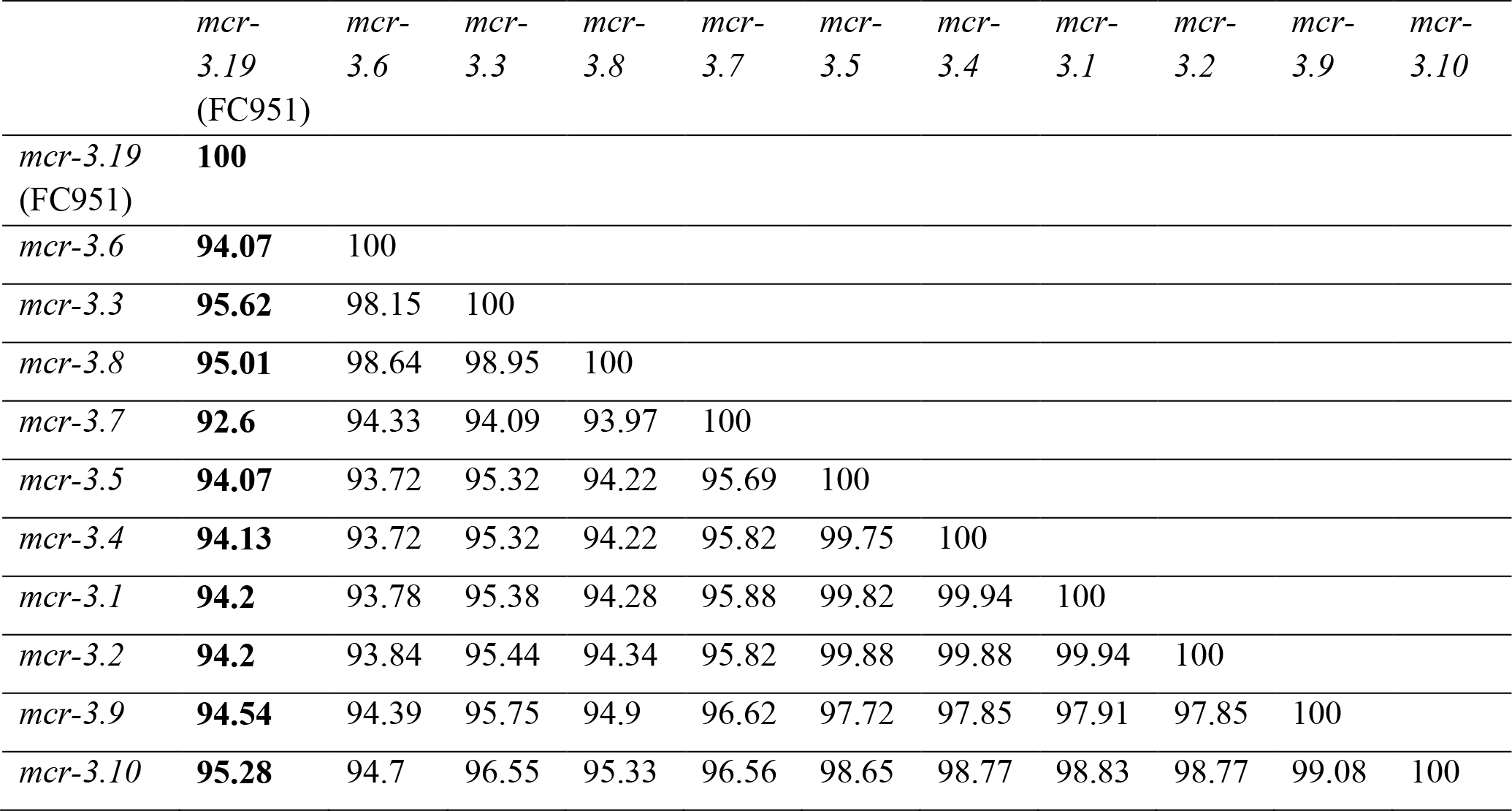
Percentage identity matrix of *mcr-3* gene variants in comparison with FC951 *mcr-3.19*

### *In silico* structure analysis

The 3D structures were modeled using the sequences for the variants *mcr*-*3.1* – *mcr*-*3.10* using the Swiss-Model server. The template used and its respective PDB ID, coverage range, and coverage identity are tabulated in Table 3. Further, the quality of the modeled variants was evaluated using the Rampage server. The percentage of the amino acids in the favored region, the percentage of amino acids in allowed region and percentage of amino acids in outlier region are tabulated in Table 3. Superimpose structural evaluation of MCR variant 3.19 against the closely related MCR variant 3.3 and MCR variant 3.10 was visualized using PyMOL (Figure 3). In addition, LEU53, ILE164, ALA57, TYR175, GLN186, ILE189, VAL176, GLY179, ALA192, PHE32, LEU50, VAL61, ARG180, VAL178, ASN182, LEU185, PHE46, LEU36, SER183, VAL29, GLY28, PRO51, LEU54, ASN25, ALA21, TRP26, GLU188, LEU24, LEU58, PRO191, ASN193, VAL195, PHE65, and ASN196 were identified as the active site of the MCR-3.19 variant using Meta Pocket server (Figure 4).

**Table 3:** List of parameters considered for modeling the variants using Swiss Model and Ramachandran plot evaluation of the structures using RAMPAGE.

| **Variants**<br>**of *mcr*** | **Template**<br>**PDB** | **Coverage**<br>**Range** | **Coverage** | **Identity** | **% of amino**<br>**acids in the**<br>**favored**<br>**region** | **% of amino**<br>**acids in the**<br>**allowed**<br>**region** | **% of amino**<br>**acids in**<br>**outlier**<br>**region** |
| --- | --- | --- | --- | --- | --- | --- | --- |
| 3.1 | 5FGN | 6-544 | 0.98 | 37.55 | 94.2% | 4.9% | 0.9% |
| 3.2 | 5FGN | 6-545 | 0.98 | 37.55 | 94.2% | 4.9% | 0.9% |
| 3.3 | 5FGN | 1-546 | 0.99 | 35.82 | 94.0% | 5.1% | 0.9% |
| 3.4 | 5FGN | 6-544 | 0.98 | 37.55 | 94.4% | 4.7% | 0.9% |
| 3.5 | 5FGN | 6-546 | 0.97 | 40.27 | 95.9% | 3.6% | 0.6% |
| 3.6 | 5FGN | 6-546 | 0.97 | 39.69 | 95.7% | 3.6% | 0.8% |
| 3.7 | 5FGN | 6-546 | 0.97 | 40.08 | 95.1% | 4.1% | 0.8% |
| 3.8 | 5FGN | 6-546 | 0.97 | 39.69 | 95.7% | 3.6% | 80.0% |
| 3.9 | 5FGN | 6-546 | 0.97 | 40.08 | 95.3% | 4.0% | 80.0% |
| 3.10 | 5FGN | 6-544 | 0.97 | 37.74 | 94.6% | 4.5% | 90.0% |
| 3.19 | 5FGN | 6-469 | 0.98 | 36.24 | 93.7% | 4.6% | 1.7% |

**Figure 3.**
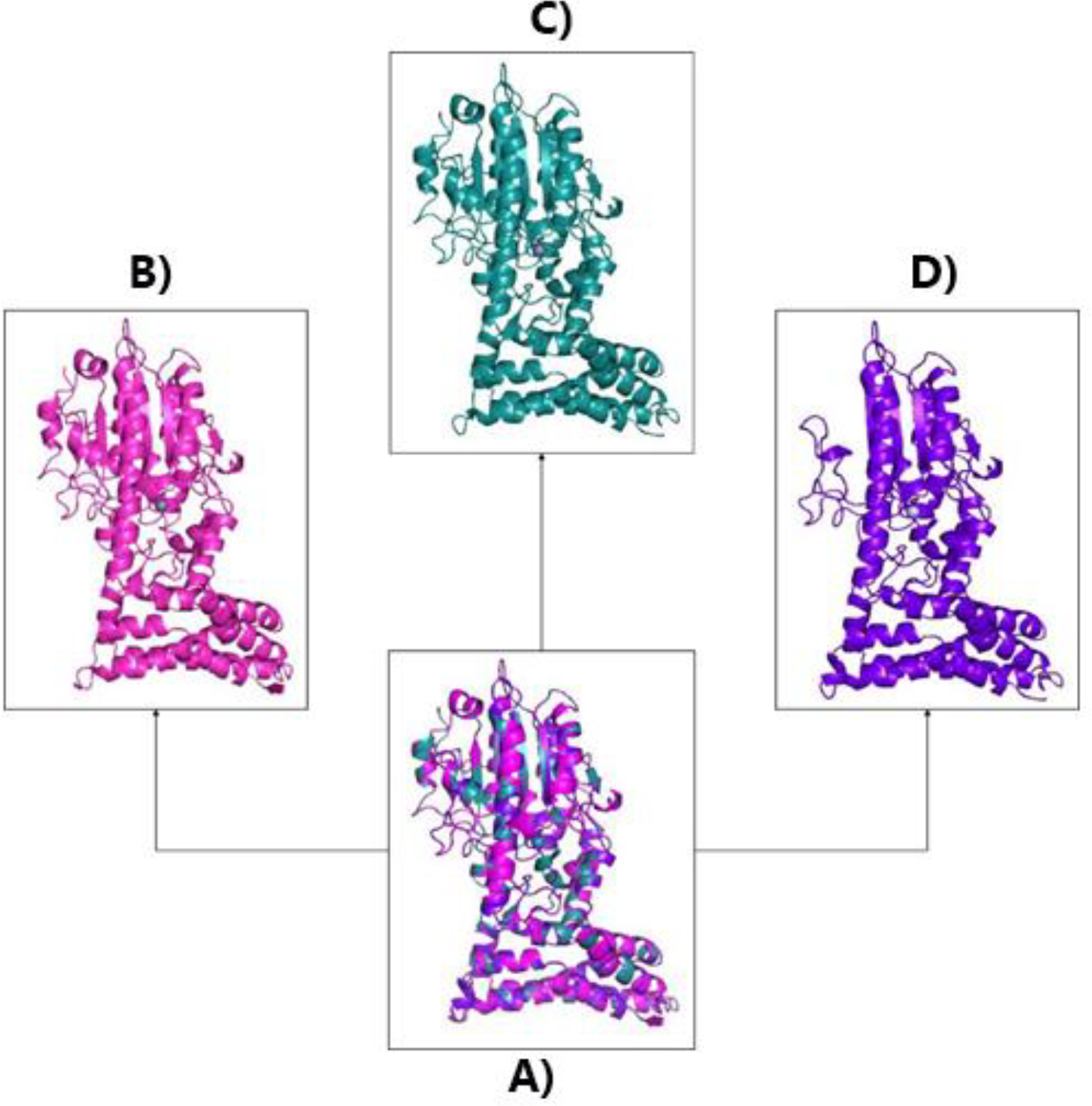
Superimpose structural evaluation of MCR variant 3.19 against the closely related MCR variant 3.3 and MCR variant 3.10. (A) Superimpose structural evaluation of MCR variant 3.19 against the closely related MCR variant 3.3 and MCR variant 3.10, (B) 3D structure of MCR variant 3.3, (C) 3D structure of MCR variant 3.10, and (D) 3D structure of MCR variant 3.19.

**Figure 4:**
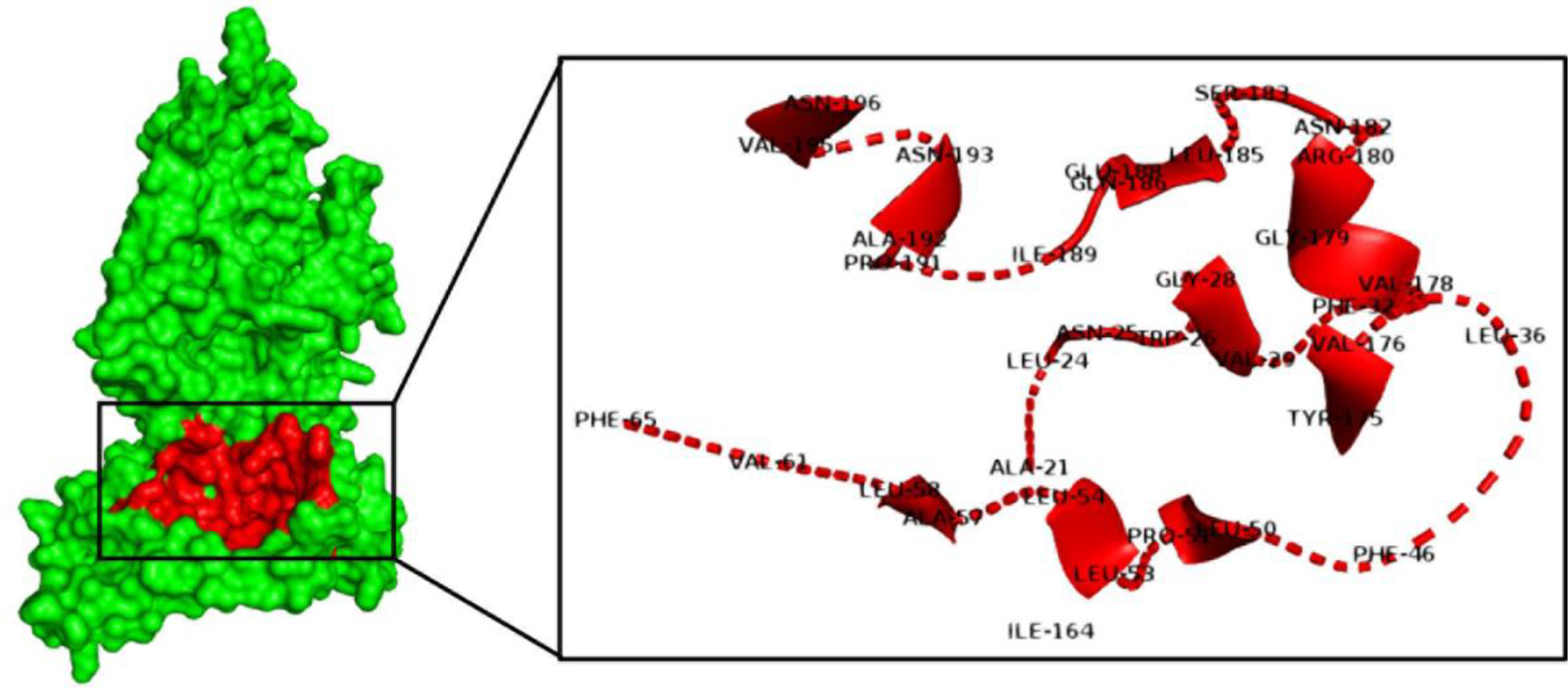
LEU53, ILE164, ALA57, TYR175, GLN186, ILE189, VAL176, GLY179, ALA192, PHE32, LEU50, VAL61, ARG180, VAL178, ASN182, LEU185, PHE46, LEU36, SER183, VAL29, GLY28, PRO51, LEU54, ASN25, ALA21, TRP26, GLU188, LEU24, LEU58, PRO191, ASN193, VAL195, PHE65, and ASN196 amino acids were identified as active sites of MCR-3.19 using Meta Pocket server.

## Discussion

Resistance to colistin is mediated mainly via alteration in the lipopolysaccharides (LPS) of the bacterial outer membrane. The alterations include mutations in lipid A modifying genes. The most commonly reported mutations were in *mgrB* gene and therefore not transferable through horizontal gene transfer [5]. However, in 2015, the first plasmid-mediated colistin resistance gene (*mcr-1*) was reported [6], which belong to the phosphoethanolamine transferase enzyme family (*Ept*A). *mcr-1* was identified in *Escherichia coli* from human patients and animals in China. In 2016, another study reported the mobilizable colistin resistance gene, *mcr-2* was from porcine and bovine *E. coli* isolates in Belgium [7]. Also, *mcr-3* and *mcr-4* genes were identified in *E. coli, Klebsiella spp* and *Salmonella spp* [4, 8]. Recently, *the mcr-5* gene was identified in *Salmonella enterica* subsp. enterica serovar Paratyphi B isolated from poultry, Denmark [9].

Considering that several mobile colistin resistance genes were identified, only *mcr-1* and *mcr-3* were reported with a high number of variants in GenBank database. A recent study highlighted the importance of third mobile colistin resistant gene, *mcr-3* in *Aeromonas salmonicida* due to its resemblance to various other phosphoethanolamine transferases in Enterobacteriaceae and also suggests that this resistance gene might have already been widely disseminated [4]. Here we discuss the novel variant of *mcr-3* identified in *Aeromonas veronii* isolated from the clinical specimen.

Novel *mcr-3* variant identified in this study, exhibited ≤95% nucleotide sequence similarity to all other previously reported *mcr*-3 variants, henceforth named *mcr-3.19*. *In silico* protein sequence comparison revealed the novelty of MCR-3.19. The superimposed protein structure comparison with MCR-3.3 and MCR-3.10 further confirmed the MCR-3.19 variant. Major structural changes were observed in domain 2. Similar comparisons of MCR-3 and MCR-1 protein structures were previously reported by Yin et al. 2017.

*mcr-3* was first identified in a 261-kb IncHI2-type plasmid pWJ1 from *E. coli* [4]. However, initially, known plasmid replicons were not identified in FC951 harboring *mcr-3.19* via PlasmidFinder. Later, on sequence assembly and alignment with pWJ1 revealed an Inc-W like replicase gene in the extrachromosomal region along with TraI and TrbN genes responsible for multi-functional conjugation and conjugal transfer proteins.

There had been a previous report on chromosome integration of *mcr-3* variant in *A. veronii* [10]. In this study, a major insertion of ISAs*18* with 1141 bp size belonging to the IS*4* family insertional elements was found within the *mcr-3.19* gene. ISAs*18* was previously reported only in *A. salmonicida* as a transposase [11]. The entire *mcr-3.19* region including the insertion was flanked by *eamA* and *dgkA*. These genes encode a metabolite transporter and a diacylglycerol kinase, respectively [12]. The *mcr-3.19* genetic environment also had other insertional elements such as ISAs*19*, ISAs*20*, IS*630*, ISKpn*15*, IS*5* and IS*3* transposase.

However, *mcr-3.19* gene disruption caused due to the insertion of ISAs*18* has rendered it non-expressive. Accordingly, the isolate FC951, in spite of harboring *mcr-3.19* gene was susceptible to colistin. In contrast, isolates resistant to colistin by MIC was negative for screened *mcr* genes, this could be due to other chromosomal mechanisms which needs to be explored.

Apart from colistin resistant gene, the genome analysis revealed the presence of other resistance genes such as, *bla*OXA-12 belong to class D beta-lactamase in FC951 conferring resistance to ampicillin, which was known to be naturally produced by *Aeromonas jandaei* and has strong activity against oxacillin [13]. The isolate also harbored *bla*CEPH-A3, which was the most common metallo-beta-lactamase (MBL) produced by *Aeromonas* species responsible for carbapenem resistance.

## Conclusion

To the best of our knowledge, this is the first report on novel *mcr-3* variant (*mcr-3.19*) at the structural level, in comparison with the known variants (MCR-3.3 – MCR-3.10). *mcr-3.19* identified in this study is non-functional for colistin resistance due to insertion of ISAs*18* within the gene. Besides, this is the first complete genome sequence of *A. veronii* from India, and hybrid genome of *A. veronii* globally. These findings indicates an extended screening of known and further exploration of unknown colistin resistance mechanisms in this pathogen as well as in other Gram-negative pathogens.

## Materials and Methods

### Isolates and identification

A total of, 30 *Aeromonas spp* isolated from stool specimen collected during January to December 2017 from the symptomatic patients attending Christian Medical College, Vellore were included in the study. Isolation and identification of the genus and species was carried out using a standard culture and biochemical tests [14].

### Antimicrobial susceptibility testing (AST)

#### Disc diffusion

AST testing was carried out using the Kirby-Bauer disk diffusion method. The antimicrobial agents tested were trimethoprim-sulfamethoxazole (1.25/23.75 µg), tetracycline (30 µg), ciprofloxacin (5 µg), cefotaxime (30 µg), imipenem (10 µg) and meropenem (10 µg) (Oxoid, UK). Quality control strains (*K. pneumoniae* ATCC 700603, *P. aeruginosa* ATCC 27853 and *E. coli* ATCC 25922) were included in all batches as recommended by the Clinical and Laboratory Standards Institute (CLSI-M45) [15].

#### Minimum Inhibitory Concentration (MIC)

Colistin MIC was determined for the studied isolates by broth microdilution and interpreted using CLSI 2017 breakpoint recommendation [16]. *mcr-1* positive *E. coli* with the expected range 4 – 8 µg/ml, *E. coli* ATCC 25922 (0.25 – 2 µg/ml) and *P. aeruginosa* ATCC 27853 (0.5 – 4 µg/ml) were used as quality control (QC) strains for colistin MIC determination.

### Screening of *mcr* genes by PCR

The presence of *mcr-1, mcr-2, mcr-3* and *mcr-4* genes encoding for plasmid-mediated colistin resistance was screened by PCR as described earlier [4, 6, 7 and 8].

### Next-generation sequencing

The isolate positive for *mcr* gene was selected for next-generation sequencing to analyze colistin resistance determinants and other genetic factors. QIAamp DNA Mini Kit (QIAGEN, Hilden, Germany) was used for genomic DNA extraction. Whole genome sequencing (WGS) of the isolate was performed with 400-bp read chemistry using an IonTorrent^TM^ Personal Genome Machine^TM^ (PGM) (Life Technologies, Carlsbad, CA) as per manufacturer’s instructions. Data were assembled de-novo using Assembler SPAdes v.5.0.0.0 embedded in Torrent Suite Server v.5.0.3. Sequence annotation was performed using PATRIC, the bacterial bioinformatics database and analysis resource (http://www.patricbrc.org), and NCBI Prokaryotic Genomes Automatic Annotation Pipeline (PGAAP, http://www.ncbi.nlm.nih.gov/genomes/static/Pipeline.html).

The CGE server (http://www.cbs.dtu.dk/services) and PATRIC [17] were employed for downstream analysis. ResFinder 2.1 (https://cge.cbs.dtu.dk//services/ResFinder/) was used for analyzing the resistance gene profile [18]. Antimicrobial resistance genes were also screened using the Antibiotic Resistance Genes Database (ARDB) and Comprehensive Antibiotic Resistance Database (CARD) through PATRIC. PlasmidFinder 1.3 (https://cge.cbs.dtu.dk//services/PlasmidFinder/) was used to screen for the presence of plasmids [19].

MLST 1.8 (MultiLocus Sequence Typing) tool was employed for sequence type analysis (https://cge.cbs.dtu.dk//services/MLST/) [20]. The genome was screened for insertion sequence elements using ISFinder (https://www-is.biotoul.fr/blast.php) [21].

### MinION Oxford Nanopore sequencing

DNA library preparation and sequencing was prepared using SQK-LSK108 Kit R9 version (Oxford Nanopore Technologies, Oxford, UK) using 1D sequencing method according to manufacturer’s protocol. Sequencing was performed using FLO-MIN106 R9 flow cell in MinION Mk 1B sequencer. MinKNOW software ver. 1.15.1 (Oxford Nanopore Technologies, Oxford, UK) was employed in a Windows platform to perform sequencing and raw data (fast5 files) were obtained.

### MinION sequence analysis

The Fast5 files were generated from MinION sequencing and the reads were basecalled with Albacore 2.0.1 (https://nanoporetech.com/about-us/news/new-basecaller-now-performs-raw-basecalling-improved-sequencing-accuracy). Furthermore, the adapters were trimmed off using Porechop (https://github.com/rrwick/Porechop). Canu 1.7 [22] was used for MinION error correction and assembly with genome size of 5.0 m as input. After, denovo assembly, the contigs were polished with Nanopolish 0.10.1 (https://github.com/jts/nanopolish).

### Hybrid assembly using IonTorrent and MinION reads

To increase the accuracy and completeness of genome, a hybrid assembly using both IonTorrent and MinION reads with Unicycler (v0.4.6) was performed [23]. By default, Unicycler utilizes SPAdes [24] to assemble the short reads with different k-mers and filter out the low depth regions. Subsequently, it trims and generates the short read assembly graph. In addition, it uses Miniasm [25] and Racon [26] to assemble the MinION long reads and further the reads were bridged to determine all the genome repeats and produces complete genome assembly. Further, the short reads were polished with multiple rounds of Pilon [27] to reduce the base level errors. After assembly, the assembly statistics and Average nucleotide identity of different assemblies were evaluated using Quast [28] and OrthoANI 0.93.1 tool [29] respectively.

### *In silico* sequence analysis

The *mcr* nucleotide sequences obtained from the experimental techniques were translated to protein sequence using the online Expasy Translate Tool (https://web.expasy.org/translate/). The percentage identity of known MCR variants in comparison with novel protein was identified using BLAST search. The conserved amino acids region from the closely related variants were determined using the Clustal Omega.

### *In silico* structure analysis

The sequences of MCR variant was used to model the 3D structure of the proteins. The 3D structures of the variants were modeled using Swiss-Model [30]. The translated variant sequences were given as the input for the 3D variant modeling. Rampage server was used to evaluate the quality of the modeled variants [31]. Finally, the MetaPocket server was used to predict the active pocket of the novel variant [32]. The structure visualization was done using PyMOL.

## Declarations

### Ethics approval and consent to participate

Not applicable

### Consent for publication

Not required

### Availability of data and material

The genome was submitted to GenBank under accession number CP032839 (Plasmid accession number CP032840).

## Authors’ contributions

SA, BV, NKDR and DPMS designed the study. DPMS and NKDR analysed and interpreted data and wrote the manuscript. DM, KA and RGNM carried out bench work and generated data. KV performed the hybrid genome assembly. TKD and GPDC performed the bioinformatics part. SA and BV reviewed and approved the manuscript. All authors have read and approved the final version of this manuscript.

## Acknowledgements

The authors thank the institutional review board for approving the study (IRB min no.8216/dated 27-02-2013).

